# Targeted anti-BCR-ABL+ ALL therapy may benefit the heart

**DOI:** 10.1101/2021.06.04.447070

**Authors:** Hanna Kirchhoff, Melanie Ricke-Hoch, Katharina Wohlan, Stefan Pietzsch, Ümran Karsli, Sergej Erschow, Robert Zweigerdt, Arnold Ganser, Matthias Eder, Michaela Scherr, Denise Hilfiker-Kleiner

**Author notes:** **Corresponding author:** Melanie Ricke-Hoch, Department of Cardiology and Angiology, Hannover Medical School, 30625 Hannover, Germany. equally contributing first and last authors.

## Abstract

Targeted therapies are currently considered the best cost-benefit anti-cancer treatment. In hematological malignancies, however, relapse rates and non-hematopoietic side effects including cardiotoxicity remain high. We here describe significant heart damage due to advanced acute lymphoblastic leukemia with t(9;22) encoding the bcr-abl oncogene (BCR-ABL+ ALL) in murine xenotransplantation models. Echocardiography reveals severe cardiac dysfunction with impaired left ventricular function and reduced heart and cardiomyocyte dimensions associated with increased apoptosis. This cardiac damage is fully reversible, but cardiac recovery depends on the therapy used to induce ALL remission. Chemotherapy-free therapy with dasatinib and venetoclax (targeting the BCR-ABL oncoprotein and mitochondrial Bcl2, respectively), as well as dexamethasone can fully revert cardiac defects whereas depletion of otherwise identical ALL in a genetic model using HSV-TK cannot. Mechanistically, dexamethasone induces pro-apoptotic BIM expression and apoptosis in ALL cells but enhances pro-survival BCLXL expression in cardiomyocytes and clinical recovery with reversion of cardiac atrophy. These data demonstrate that therapies designed to optimize apoptosis induction in ALL may circumvent cardiac on-target side effects and may even activate cardiac recovery. In the future, combining careful clinical monitoring of cardiotoxicity in leukemic patients with further characterization of organ-specific side effects and signaling pathways activated by malignancy and/or anti-tumor therapies seems reasonable.

## Main text

Targeted therapies are currently considered the best cost-benefit anti-cancer treatment. Accordingly, mechanism-based interventions can rationally be designed and evaluated in murine xenograft models with regard to clinically relevant end points (1, 2). However, targeted anti-tumor therapies may exert unexpected on- and off-target side effects in other organ systems not affected by malignant transformation or malignant cells. For example, cardiovascular toxicities have been described for trastuzumab, bevacizumab, imatinib and sunitinib in solid tumors and in chronic myeloid leukemia (CML), respectively (3). We therefore hypothesized that murine xenograft models may be used to both optimize anti-leukemic therapy and to characterize and prevent side effects on the heart where applicable.

We recently described a curative combined pharmacotherapy with dasatinib (DAS), venetoclax (VEN) and dexamethasone (DEX) in a humanized mouse model for BCR-ABL+ ALL (BV173-mice: BV173 ALL cell line with stable luciferase expression for bioluminescence in vivo imaging (BLI) to visualize leukemic burden in NSG mice) (1). In the present study, BV173-mice developed aggressive leukemia with a mean survival of 32±2 days. At advanced stage of leukemia (around week 4 after cell injection) mice displayed severe cardiac dysfunction as assessed by echocardiography in comparison to age-matched healthy controls (Figure 1A, Table 1). This cardiac phenotype included significantly reduced heart size and substantially impaired left ventricular (LV) function due to reduced cardiac output (LV CO) and stroke volume (ESV) (Figure 1B, C, D, Table 1). Post mortem analysis revealed reduced heart weight with no evidence of infiltrating leukemic cells in LV tissue (Table 1 and data not shown). Together with the reduced total body weight (Table 1) this phenotype suggests cachexia and cardiac atrophy in mice with advanced leukemia. Accordingly, we observed a reduction of cardiomyocyte (CM) cross-sectional area (CSA) (Figure 1 E, F) associated with enhanced cardiac expression of *microtubule-associated light chain 3 beta* (*LC3B*) (Figure 1G) and BCL2/adenovirus E1B 19 kDa protein-interacting protein 3 (BNIP3) both linked to cardiac atrophy in leukemic as compared to control mice (Figure 1H, I) (4-6). Beyond cardiac atrophy, TUNEL assays revealed an increase in rare cardiomyocyte apoptosis in leukemic compared to control mice with no evidence of elevated cardiac fibrosis or inflammation (Figure 1J, Supplemental Figure 1A and data not shown). Almost identical data were found with an independent patient-derived xenograft (PDX) BCR-ABL+ ALL model (Supplemental Figure 1 B-I, Table 1).

**Figure 1.**
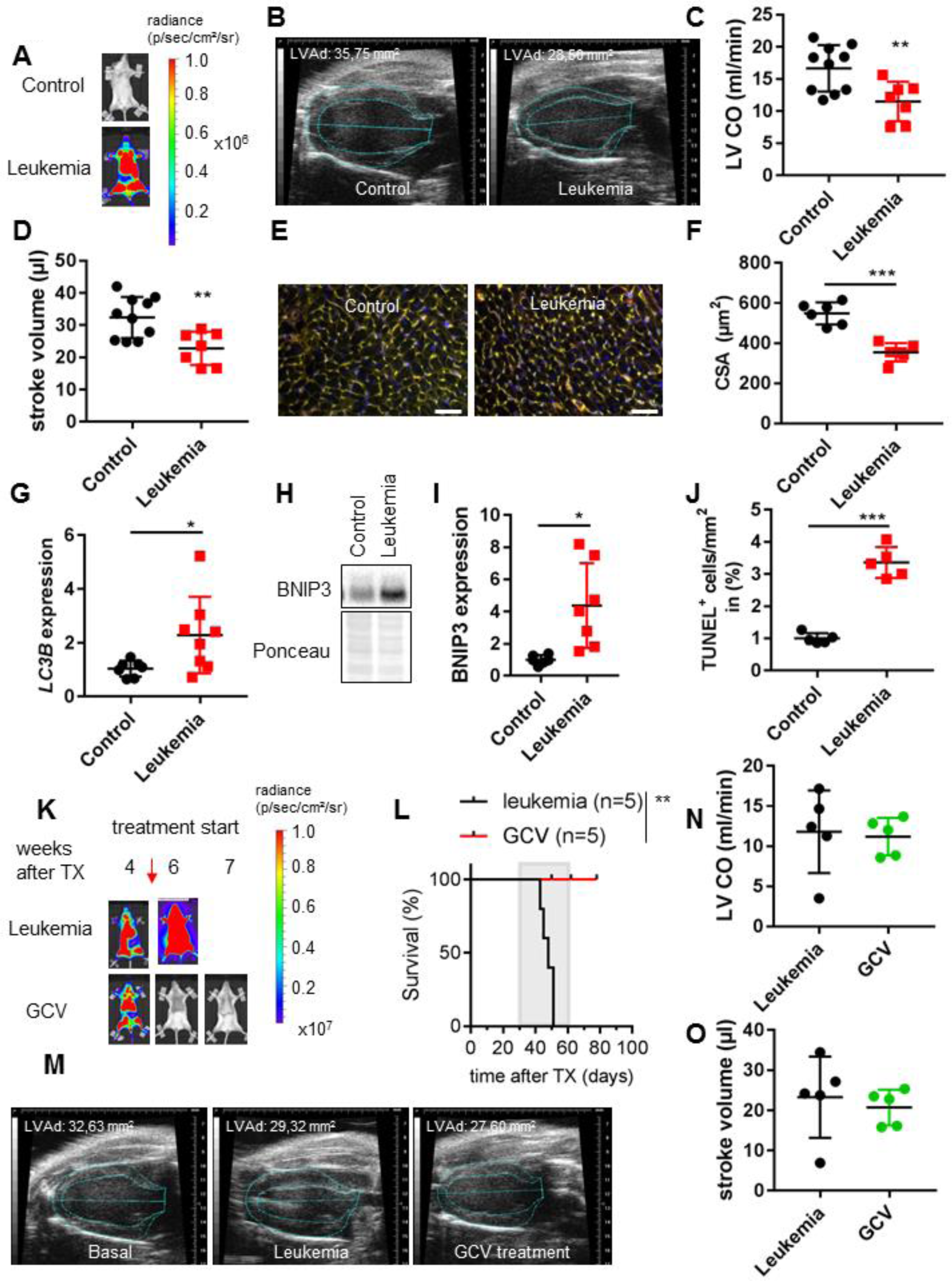
Cardiac phenotype of BCR-ABL^+^ ALL. **(A)** Representative BLI of NSG mice non-transplanted (control) or transplanted with luciferase-expressing BV173 cells 4 weeks after inoculation. **(B)** Representative echocardiographic picture in parasternal long axis view at end-diastole of BV173 and control heart at severe state of disease 4 weeks after transplantation. Left ventricular area diastole (LVAd) indicates cardiac dimensions and size. **(C**-**D)** Left ventricular cardiac output (LV CO) (C) and stroke volume (D) of age matched healthy controls (control) or BV173-mice at severe state of disease 4 weeks after transplantation. **p<0.01. **(E)** LV cryosections stained with isolectin B4 (blood vessels, green), wheat germ agglutinin (WGA, cell membranes, red) and nuclei (DAPI, blue), scale bar: 50 μm. **(F)** Dot plot summarizes cardiomyocyte cross sectional area (CSA) in BV173 (n=6) and control (n=5) LVs at severe state of disease 4 weeks after transplantation. ***p<0.001. **(G)** LC3B mRNA expression *p<0.05. **(H-I)** Representative immunoblot and densitometric quantification (n=7) of LV tissue for BNIP3. Ponceau served as loading control. *p<0.05. **(J)** Dot plot summarize the number of TUNEL positive nuclei per mm^2^ in BV173 (n=5) and control (n=5) LVs at severe state of disease 4 weeks after transplantation. ***p<0.001. **(K)** Serial BLI of NSG mice transplanted with luciferase and herpes simplex thymidin kinase expressing BV173 cells. Mice were monitored up to 7 weeks after inoculation. GCV treatment started 4 weeks after transplantation. **(L)** Kaplan Meier survival curve for untreated (leukemia n=5) and GCV treated (n=5) BV173-mice. Grey area indicates the GCV treatment period. Log-rank test was used for statistical survival analysis. **p<0.01. **(M)** Representative serial echocardiographic pictures in parasternal long axis view at end-diastole of NSG hearts at basal state, severe leukemia disease state and after three weeks of GCV treatment. Heart size shown as LVAd. **(N-O)** Left ventricular cardiac output (LV CO) (N) and stroke volume (O) of BV173-mice at severe state of disease or after three weeks of GCV treatment. Underlying data can be found in Figure 1-source data folder.

**Table 1.** Morphometry and cardiac function of NSG mice transplanted with BCR-ABL+ ALL. Body weight (BW), heart weight (HW), left ventricular end-diastolic diameter (LVEDD), left ventricular end-systolic diameter (LVESD), end-diastolic area (EDA), end-systolic area (ESA), heart rate (HR), endocardial diastolic volume (EVd), endocardial systolic volume (EVs), fractional area change (FAC), ejection fraction (EF) determined in NSG mice (cell lines BV173 and PDX). *p<0.05, **p<0.01, ***p<0.001 students t-test to corresponding aged-matched healthy control group.

|  | Control (n=10) | BV173 (n=7) | Control (n=9) | PDX (n=5) |
| --- | --- | --- | --- | --- |
| BW [g] | 24.12±1.68 | 18.53±1.85* | 28.06±1.01 | 22.10±1.81*** |
| HW [mg] | 98.46±0.88 | 68.46±8.29*** | 102.46±6.83 | 86.20±5.07** |
| HW/BW [mg/g] | 4.29±0.55 | 3.7±0.47 | 3.65±0.15 | 3.92±0.39 |
| LVEDD [mm] | 7.71±0.64 | 6.65±0.36** | 7.56±0.37 | 7.19±0.78 |
| LVESD [mm] | 6.35±0.51 | 5.33±0.68 | 6.48±0.45 | 6.41±0.31 |
| EDA [mm<sup>2</sup>] | 0.20±0.02 | 0.16±0.015** | 0.20±0.01 | 0.16±0.03* |
| ESA [mm<sup>2</sup>] | 0.09±0.02 | 0.07±0.02 | 0.11±0.02 | 0.10±0.02 |
| HR [bpm] | 511±30 | 499±34 | 506±37 | 492±36 |
| EVd [μl] | 43.87±7.75 | 31.90±5.58* | 43.44±5.39 | 32.39±9.84* |
| EVs [μl] | 11.36±3.24 | 9.09±4.33 | 15.75±3.80 | 12.58±4.60 |
| FAC [%] | 54.36±5.56 | 53.28±11.56 | 44.80±7.80 | 41.60±7.96 |
| EF [%] | 74.14±5.58 | 71.84±11.96 | 63.66±7.80 | 61.51±9.72 |

We next used a genetic model to analyze leukemia-dependency of this cardiac phenotype. NSG mice were injected with BV173 cells expressing herpes simplex virus-thymidine kinase (HSV-TK) for selective ablation of transduced cells upon treatment with ganciclovir (GCV) (7). Upon advanced leukemic engraftment at 4 weeks, mice were treated intraperitoneally with GCV leading to rapid depletion of leukemia and prolonged survival (Figure 1 K, L). However, no significant improvement of ALL-induced cardiac failure was detected three weeks after remission induction (Figure 1 M, Supplement Table 1 - Supplementary file 1) with no recovery of cardiac dimensions (decline in LV area, diastole (LVAd) and cardiac dysfunction (impaired LV CO and ESV) at the end of GCV therapy (Figure 1 M, N, O). Cardiotoxicity of GCV therapy could be excluded by treatment of healthy mice with GCV for 4 weeks with no detectable cardiotoxicity (Supplemental Table 2 – Supplementary file 1). These data demonstrate severe cardiac damage at late stages of ALL progression, which is not reverted by remission of leukemia.

We next treated BV173-mice with advanced leukemia (Figure 2A) using either the DAS/VEN/DEX triple- or the DAS/VEN double-therapy for 4 weeks and analyzed leukemic burden, survival and cardiac parameters. Therapy with DAS/VEN/DEX but not DAS/VEN led to leukemic regression and prolonged survival beyond the course of treatment (Figure 2B). In contrast to GCV treatment, therapy with DAS/VEN/DEX was associated with improved cardiac dimensions, heart weight and LV function (Figure 2C and Supplemental Table 3 - Supplementary file 1). In addition, CSA was preserved (Figure 2D, E) and no induction of apoptosis was detected (Figure 2F) with no difference in cardiac phenotype compared to age-matched healthy controls. In an experiment closer to the situation seen in the clinics for ALL patients, BV173-mice were treated with the DAS/VEN/DEX triple combination at an earlier leukemic stage (around one week after cell injection) and cardiac and morphometric analyses were performed in a follow-up cohort of long-term survivors after 38 weeks. No signs of cardiac dysfunction or alterations in cardiac and CM morphometric dimensions (Supplemental Table 4 – Supplementary file 1) were detected without any differences to age-matched healthy controls (Supplemental Figure 2). These data clearly demonstrate that cardiac outcome depends on the kind of anti-leukemic therapy in this model.

**Figure 2.**
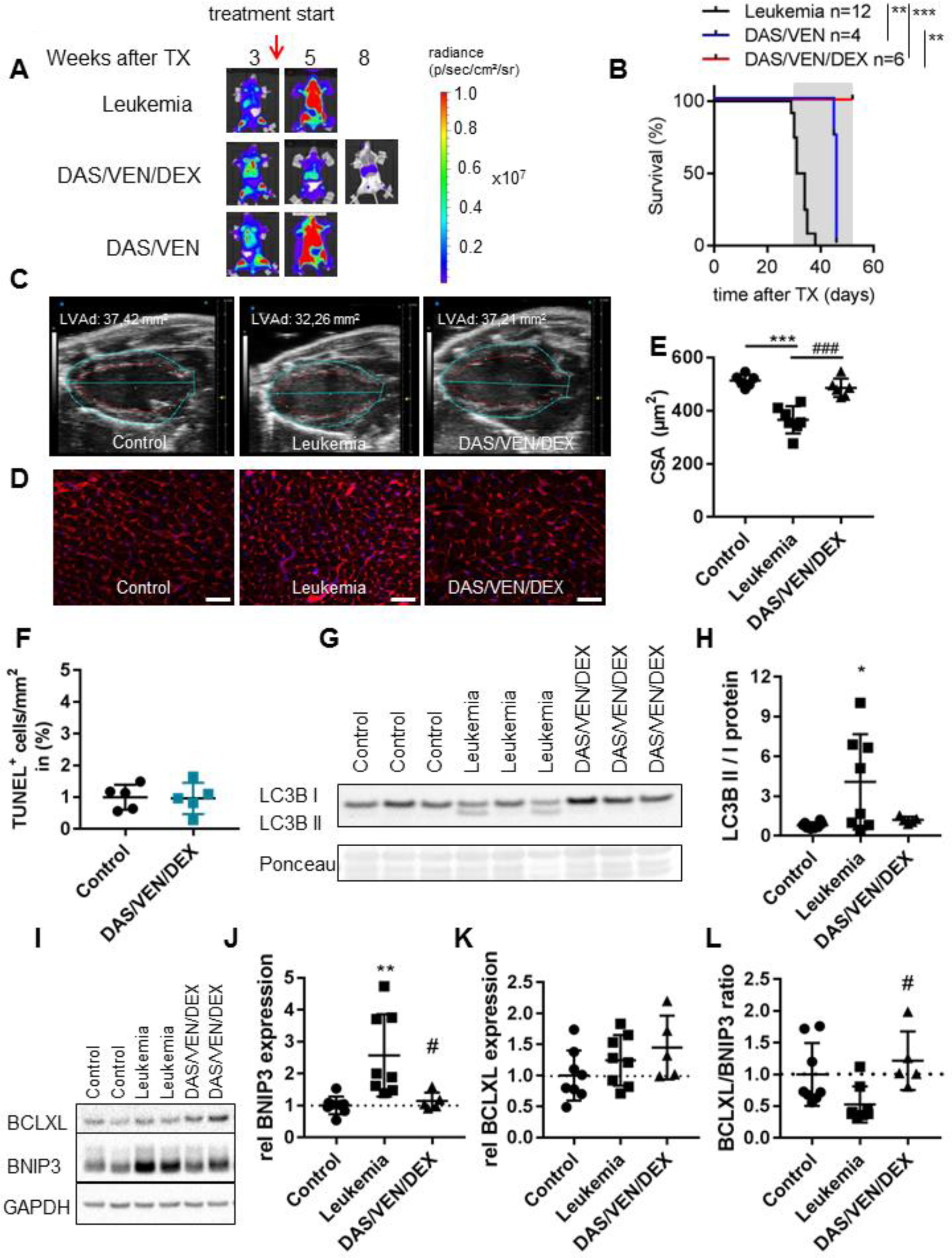
Treatment at advanced state of disease can restore cardiac function. **(A)** Representative serial BLI analysis of NSG mice transplanted with luciferase expressing BV173 cells. Mice were treated at advanced state of leukemia 4 weeks after transplantation with DAS/VEN/DEX (n=6) or DAS/VEN (n=4). **(B)** Kaplan Meier survival curve for BV173-mice treated with DAS/VEN/DEX (n=6) or DAS/VEN (n=4). Grey area indicates the treatment period. Log-rank test was used for statistical survival analysis. **p<0.01, ***p<0.001. **(C)** Representative echocardiographic picture in parasternal long axis view at end-diastole of age matched healthy control, untreated BV173 and DAS/VEN/DEX treated BV173 hearts at 4 weeks after treatment start. LVAd indicate cardiac dimensions and size. **(D)** Representative LV cryosections stained with WGA (cell membranes, red) and nuclei (DAPI, blue), scale bar: 50 μm. **(E)** Dot plot summarizes cardiomyocyte CSA of healthy control (n=6), untreated BV173 (n=7) and DAS/VEN/DEX treated BV173 (n=6) hearts at 4 weeks after treatment start. ***p<0.001 vs. Control, ^###^p<0.001 vs Leukemia. **(F)** Dot plot summarize the number of TUNEL positive nuclei per mm^2^ in healthy control (n=5) and DAS/VEN/DEX treated BV173 (n=5) hearts, not significant. **(G-H)** Immunoblot analysis and densitometric quantification of LV tissue of healthy (control) or BV173-mice treated with vehicle or DAS/VEN/DEX. Expression of LC3B isoforms was analysed. Ponceau served as loading control. *p<0.05. **(I)** Immunoblot analysis of LV tissue of healthy (control) or BV173-mice treated with vehicle or DAS/VEN/DEX. Expression of BCLXL and BNIP3 was analysed. GAPDH served as loading control. **(J-L)** Densitometric quantification of BNIP3 (J), and BCLXL (K) and ratio of BCLXL/BNIP3 (L) expression. *p<0.05 vs. Control, #p<0.05 vs. Leukemia. Underlying data can be found in Figure 2-source data folder.

The DAS/VEN/DEX triple combination has been optimized for apoptosis induction in ALL cells based on increased mitochondrial outer membrane permeabilization (MOMP) (1, 2). In these cells DEX and DAS enhance BIM loading to BCL2 thereby sensitizing for VEN cytotoxicity (1, 2). Surprisingly and in contrast to its effect on ALL cells the DAS/VEN/DEX triple-therapy does not enhance the low level of apoptosis in the heart but rather induces cardiac recovery from ALL induced damage. BIM binding to BCL2 is the main predictor of VEN cytotoxicity in ALL cells. In CM however, BIM is silenced during postnatal development, and adult CM do not rely on BCL2 for survival (8-10). We therefore analyzed cardiac expression of additional mitochondrial apoptosis regulators BCLXL and MCL1, as well as that of BNIP3 and LC3B involved in regulation of cellular atrophy (Figure 2 G-L, Supplemental Figure 3A-C). BCLXL has a dual function as an anti-apoptotic protein and as a regulator for BNIP3 and LC3B (6, 11-13). BCLXL and MCL1 are expressed at similar levels in leukemic and control mice (Supplemental Figure 3A, B). In contrast, we found heterogeneous but significantly increased ratios of LC3B isoforms II/I in LV tissue indicating elevated numbers of autophagolysosome vesicles and increased BNIP3 levels in leukemic as compared to control mice (Figure 2G-J). BNIP3 is not expressed in leukemic BV173 cells (Supplemental Figure 3C). Upon DAS/VEN/DEX treatment both enhanced cardiac BNIP3 expression and the increased ratio of LC3B isoforms II/I were reverted (Figure 2G-J). Furthermore, cardiac BCLXL expression increased upon DAS/VEN/DEX but not DAS/VEN treatment (Figure 2K, L, Supplemental Figure 3D-E). We next analyzed RNA expression of *BNIP3* and its binding partner *BCLXL* in cell cultures of primary adult mouse CM (Supplemental Figure 3F-G) and human iPSC-derived CM (iPSC-CM) (Supplemental Figure 3H-I) treated with DAS, VEN, and DEX. Only DEX, but neither VEN nor DAS induced *BCLXL* expression in both adult and iPSC-CM. *BNIP3* expression was not affected by any drug (Supplemental Figure 3G and 3I). These data demonstrate that therapies designed to optimize apoptosis induction in ALL may circumvent cardiac on-target side effects and even activate recovery from cellular atrophy in the heart.

On the molecular level low or absent expression of BIM and BCR-ABL in LV cardiac tissue minimize the risk of on-target side effects in cardiomyocytes upon therapy with VEN and DAS, respectively. DEX however induces cardiac BCLXL expression in mice and in cardiomyocyte cell cultures and may thereby contribute to cardiac recovery from atrophy and protect from cardiomyocytic apoptosis. Although cell-, tissue-, organ-as well as species-specific effects and the underlying molecular mechanisms remain to be determined in detail these data demonstrate that murine xenograft leukemia models can be used to simultaneously optimize anti-leukemic therapies and analyze disease and/or therapy dependent cardiotoxicity. Cardiac comorbidity of current anti-tumor therapies may have tremendous impact on long-term survival and morbidity of tumor patients as most prominently described for anthracycline-induced cardiomyopathy (14). On the other side transient cardiac damage during anti-leukemic therapies as indicated by elevation of cardiac stress or ischemia markers such as NT-proBNP and troponin T as well as arrhythmias are regularly seen in patients during intensive anti-leukemic therapies (15). It therefore seems reasonable to complement careful clinical monitoring of cardiotoxicity in leukemic patients with further characterization of organ-specific side effects and signaling pathways activated by malignancy and/or anti-tumor therapies in the future.

## Material and Methods

Cell culture media were obtained from Biochrome and Invitrogen. All other chemicals except the *pharmacologic agents listed below* were purchased from Sigma-Aldrich.

### Pharmacologic agents

For *in vitro* experiments venetoclax (SelleckChem) and dasatinib (Santa Cruz Biotechnology) were solubilized in dimethyl sulfoxide (DMSO) (Merck, Darmstadt, Germany) to 10 mM stocks. For *in vivo* studies venetoclax and dasatinib were solubilized in a vehicle consisting of 60% Phosal, 30% PEG 400 and 10% Ethanol to 20 mg/mL and 40 mg/mL, respectively. Dexamethasone (SelleckChem) was solubilized in PBS at stock concentrations of 10 mM (*in vitro)* and 20 mg/mL (*in vivo*). Ganciclovir (HEXAL, Holzkirchen, Germany) was stored in stock solution of 50 mg/mL in 0.9% NaCL.

### Cell lines

The BCR-ABL-positive B-cell leukemia cell line BV173 (DSMZ, Braunschweig, Germany) was cultivated in RPMI1640 Supplemented with 10% FCS and 1% penicillin/streptomycin (Gibco).

### Preparation of lentiviral supernatants and transduction

The preparation of recombinant lentiviral supernatants and lentiviral transductions were performed as described earlier (16). Transduction efficacy was assessed by eGFP- or YFP-expression using a FACS Calibur (BD Bioscience, San Jose).

### Isolation and culture of primary adult mouse cardiomyocytes

Cardiomyocytes were isolated by enzymatic digestion from C57BL/6N as previously described and subsequently plated on laminin-coated tissue culture plates from a subset of mice (4). Isolated cardiomyocytes were cultivated in 1x MEM culture medium (Gibco) Supplemented with 1% penicillin, 1% L-glutamine, and 25 mM (-)-blebbistatin (Cayman Chem, 13013) in 1% CO_2_ at 37°C in a cell culture incubator and were treated with venetoclax (100 nM), dasatinib (150 nM) and dexamethasone (500 nM) alone or in combination for 24 hours.

### Culture and cardiomyogenic differentiation of human iPSCs

Human iPSCs were grown in mTeSR or E8 medium (STEMCELL Technologies) as described (17). In brief, cells were passaged every 3-4 days using Accutase (Life Technologies) and reseeded at 0.5×10^4^ cells/mL on Geltrex-coated (Life Technologies) culture flasks including 10 mM ROCK inhibitor (Y-27632). For the directed chemically defined differentiation, single cells were inoculated in E8 for pre-culture using Erlenmeyer flasks (20 mL working volume; rotated at 70 rpm on an orbital shaker) for aggregate formation in suspension (18). The cell density was determined after 48 h and adjusted to 5×10^5^ cells/mL for differentiation in CDM3 medium Supplemented with 5 mM CHIR and 5 mM Y-27632 as described (19). Precisely 24 h later the medium was replaced by CDM3 Supplemented with 5 mM IWP2; 48 h later the medium was replaced with pure CDM3; fresh CDM3 medium was added every 2–3 days thereafter. Resulting iPSC cardiomyocytes (iPSC-CM) where analyzed on day 10 of differentiation as reported (19) and used for experiments within 1-2 weeks. Cells were treated with venetoclax (100 nM), dasatinib (150 nM) and dexamethasone (500 nM) alone or in combination for 24 hours and total RNA was isolated using Trizol (Invitrogen).

### Animal experiments

All animal studies were in accordance with the German animal protection law and with the European Communities Council Directive 86/609/EEC and 2010/63/EU for the protection of animals used for experimental purposes. All experiments were approved by the Local Institutional Animal Care and Research Advisory Committee and permitted by the local authority, the Niedersächsisches Landesamt für Verbraucherschutz und Lebensmittelsicherheit [No. 33.14-42502-04-16/2217], [33.12.42502-04-17/2520] and [No. 33.19-42502-05-18A271].

Leukemic BV173 cells or PDX cells (1×10^6^) were transplanted intravenously into recipient female NSG (NOD.Cg-Prkdcscid Il2rgtm1WjI/SzJ) mice (10±2 weeks of age) and after cell implantation, mice received continuous analgesia (Novalgin, 1000 mg/kg/day in drinking water) and antibiosis (0.08 mg/ml in drinking water Ciprofloxacin) as described (1). The combination of DAS (10 mg/kg) and DEX was administered by oral gavage in 0.1M sodium citrate 5 days per week; venetoclax (20 mg/kg) was applied orally with a treatment delay of minimum 2 hours in a vehicle consisting of 60% Phosal, 30% PEG 400 and 10% Ethanol as described (1). The health condition of animals was assessed based on the guidelines of recognition and distress in experimental animals as proposed by Morton and Griffiths (20). Where possible, data were analyzed by an experimenter blinded to the tumor status and treatment regimen of the animals.

### In vivo bioluminescence Imaging

BV173 and PDX (L4951) cells were transduced with lentiviral vectors SLIEW (or HSVTK) encoding both enhanced GFP (or YFP), for *in vitro* analysis, and firefly luciferase, for *in vivo* bioluminescence imaging (BLI) (4, 7, 21). Tumor burden was assessed by *in vivo* bioluminescence imaging (BLI) by using an IVIS Lumina II (Caliper Life Sciences, Hopkinton, MA) (1, 2, 4). D-Luciferin (0.8 mg/mouse) (AppliChem, Darmstadt, DE) was provided intraperitoneally and whole animal imaging was performed from ventral perspective. Bioluminescence radiance was analyzed using Living Image 4.0 software.

### Echocardiography

Serial echocardiography was performed to assess heart rate (HR) and contractile function in NSG female leukemia-bearing mice (BV173 18w, PDX 28w) and in corresponding healthy age-matched controls (1-4% isoflurane inhalation, connected to a rodent ventilator) using a Vevo 770 or Vevo 3100 (Visual Sonics) as described (4). Parasternal long-axis views were recorded in B-mode at the level of the papillary muscle, and still images were used to measure LV end-diastolic diameter (LVEDD) and LV end-systolic diameter (LVESD), end-diastolic area (EDA), end-systolic area (ESA) and calculate fractional area change (FAC) ([end-diastolic area – end-systolic area / end-diastolic area x 100]). Cardiac output (LV CO), endocardial stroke volume (ESV), endocardial diastolic volume (EVd), and endocardial systolic volume (EVs) were calculated by Visual Sonics Vevo 770 software version 3 and Vevo lab 2.1.0 software.

### Histology, morphometry and immunostaining

For cardiac morphological analyses, hearts were embedded in OCT and frozen at -80°C or were fixed in formalin and embedded in paraffin as described (4). Cardiomyocyte cross-sectional area (CSA) was determined after wheat germ agglutinin (WGA)/Hoechst staining as previously described (4). For nuclear staining DAPI, Hoechst 33258 (SIGMA) staining was used.

For histological analyses of apoptosis by TUNEL assay, hearts were fixed in formalin and embedded in paraffin as described (22). Apoptotic nuclei were detected by in situ terminal deoxynucleotidyl transferase-mediated digoxigenin-conjugated dUTP nick end labeling (TUNEL) and by nuclear morphology using Hoechst 33258 staining (22). Sections were co-stained with WGA (Vector). Images were taken by fluorescence microscopy with an Axiovert 200M microscope using AxioVision 4.8 software or Axio Observer 7 using Zen 2.6 software (Carl Zeiss, Jena, Germany).

### Immunoblotting

Whole cell lysates of BV173 were prepared with lysis buffer (20 mM HEPES, pH 7.5, 0.4 M NaCl; 1 mM EDTA, 1 mM EGTA, 1 mM DTT) Supplemented with mini complete protease inhibitor cocktail tablet (Roche Diagnostics, Mannheim, Germany) (1). LV tissue was lysed as described previously (4). Protein lysates were separated by sodium dodecyl sulphate-polyacrylamide gel electrophoresis (SDS-PAGE), transferred to Hybond enhanced chemiluminescence (ECL) nitrocellulose membrane (Amersham Bioscience, Uppsala, Sweden) and membranes were incubated with the following antibodies according to the manufacturer’s protocol: anti-BCLXL (cs2762), anti-MCL1 (cs5453) and anti-GAPDH (cs2118), from Cell Signaling Technology, anti-BNIP3 from Abcam (Ab109362) and anti-LC3B from Sigma (#7543). Chemiluminescence was used for visualization using the ECL Western blotting detection reagents (PerkinElmer) according to the manufacturer. Densitometric analysis was performed using ChemiDoc MP Imaging system and ImageLab software version 5.0 (Bio-Rad).

### RNA-Isolation and qRT-PCR

Total RNA from cell lines, adult mouse CM or iPSC-derived CM was prepared using Trizol (Invitrogen), cDNA synthesis was performed with 1 μg of total RNA digested with *DNaseI* and subjected to SYBR green qRT-PCR (primer sequences are provided in Table 1) and TaqMan-based (Applied Biosystems) gene expression profiling following the manufacturer’s protocol.and as described (1, 4).

Primer/probe assays for human BCLXL (Hs00236329_m1), human BNIP3 (Hs00969291_m1) and human 18S (Hs99999901_s1) were purchased from Applied Biosystems. 18S served as an internal control. Real-time PCR was performed using an ABI7500 cycler (Applied Biosystems).

### Statistics

All data are presented as mean ±SD unless otherwise stated. P-value was calculated using two-sided students t-test or 1-way ANOVA with Bonferroni’
ss post-hoc correction for multiple group comparisons. For statistical analysis of Kaplan Meier survival curves Log-rank test was performed. P<0.05 was considered significant.

### Software

Vevo 770/3100 Imaging, Vevo lab 2.1.0 software, Image Lab software version 5, CompuSyn Software (Biosoft), Living Images 4.0 (Perkin Elmer, Waltham, MA, USA, Zeiss Software AxioVert and Zen, GraphPad Prism version 7.0 for Mac OS X (GraphPad Software, San Diego California USA).

## Acknowledgements

This work is supported by grants of H.W. & J. Hector-Stiftung to M.S. and M.E., Stiftung Gerdes to M.S. and H.K. and D. H.-K. and S.P., the Erich and Emmy Hoselmann-Stiftung to the Department of Cardiology and Angiology, the Deutsche Forschungsgemeinschaft (DFG KFO311, HI 842/10-1, HI 842/10-2 to D.H.-K. and RI 2531/2-1, RI 2531/2-2 to M.R.-H.), by REBIRTH I/II to D.H.-K., by the Foundation Leducq (Project ID 19CVD02) to D.H.-K. We thank Karin Battmer, Iris Dallmann, Birgit Brandt and Martina Kasten for excellent technical help. We acknowledge the staff of the Central Animal Facility of Hannover Medical School. We thank Olaf Heidenreich for providing us patient-derived xenograft cells L4951.

## Author information

These authors contributed equally:

First authors: Hanna Kirchhoff, Melanie Ricke-Hoch

Last authors: Matthias Eder, Michaela Scherr and Denise Hilfiker-Kleiner

## Supporting information

### Supplemental Figure Legends

**Supplemental Figure 1:**
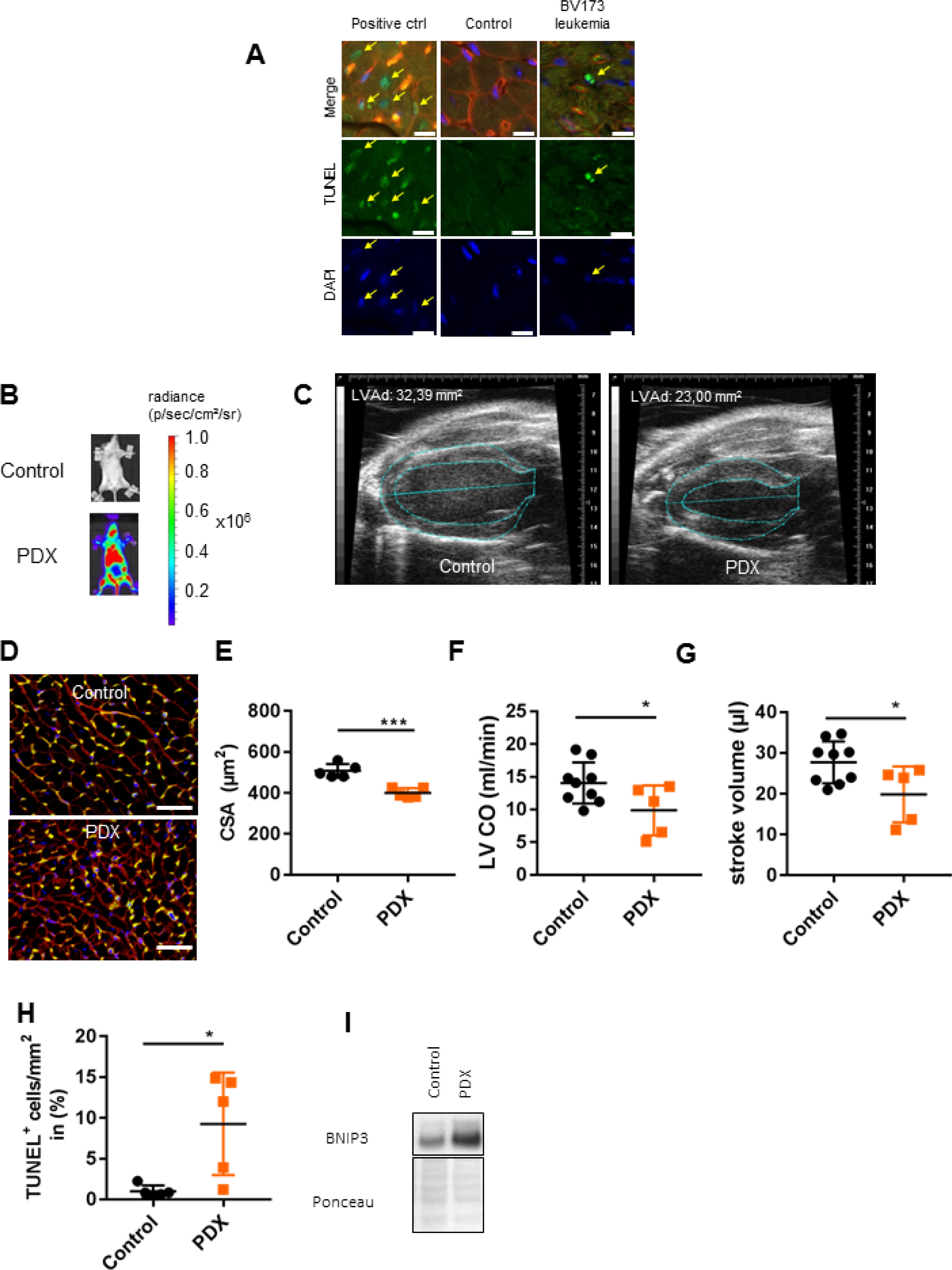
**(A)** RepresentativeTUNEL positive nuclei (green) with apoptotic morphology co-stained with WGA (red) and DAPI (blue) (upper panel: merged staining TUNEL, WGA and DAPI; middle panel: TUNEL alone; lower panel: DAPI alone) of healthy control and BV173 NSG mice. Yellow arrows indicate double positive stained cells for TUNEL and DAPI. Scale bar indicate 10 μm. **(B)** Representative BLI of NSG mice non-transplanted or transplanted with luciferase expressing PDX (L4951) cells 10 weeks after transplantation. **(C)** Representative echocardiographic picture in parasternal long axis view at end-diastole of PDX (L4951) and control hearts at severe state of disease 10 weeks after transplantation. LVAd indicates cardiac dimension and size. **(D)** Representative LV cryosections stained with isolectin B4 (blood vessels, green), WGA (cell membranes, red) and nuclei (DAPI, blue), scale bar: 50 μm. **(E)** Dot plot summarizes cardiomyocyte CSA in PDX (n=5) and control (n=5) LVs at severe state of disease 10 weeks after transplantation. ***p<0.001. **(F-G)** Left ventricular cardiac output (LV CO) (E) and stroke volume (F) of age matched healthy controls (control) (n=9) or PDX-mice (n=5) 10 weeks after transplantation. **p<0.01. **(H)** Dot plot summarize the number of TUNEL positive nuclei per mm^2^ in PDX (n=5) and healthy control (n=5), *p<0.05. **(I)** Representative Immunoblot of LV tissue of age matched healthy control mice or PDX-mice at severe state of disease for BNIP3. Ponceau served as loading control. Underlying data can be found in Supplement Figure 1-source data folder.

**Supplemental Figure 2:**
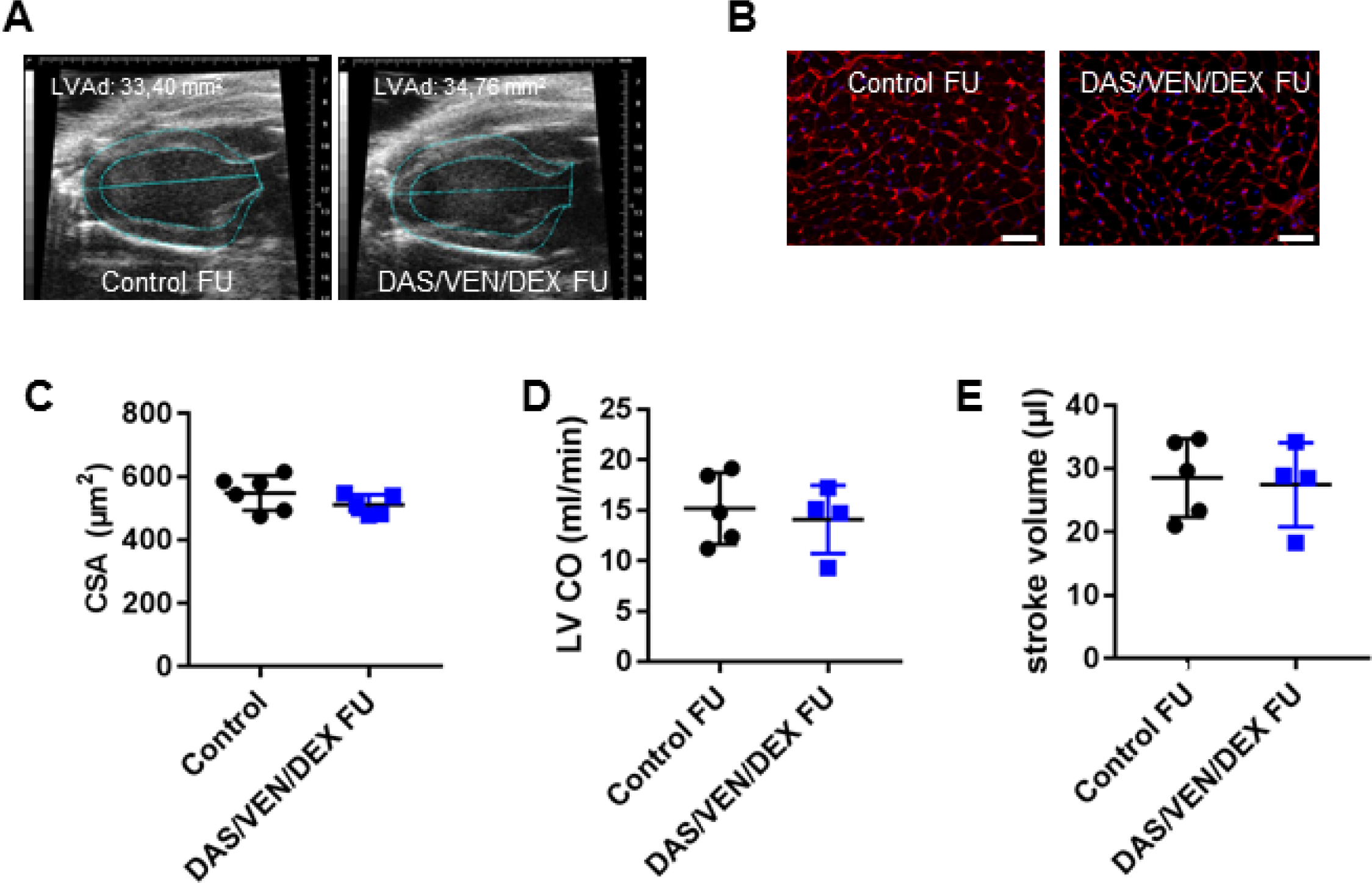
**(A)** Representative echocardiographic picture in parasternal long axis view at end-diastole of healthy control and DAS/VEN/DEX treated BV173 hearts at 35 weeks follow-up (FU) after transplantation. LVAd indicate cardiac dimensions and size. **(B)** Representative LV cryosections stained with WGA (cell membranes, red) and nuclei (DAPI, blue), scale bar: 50 μm. **(C)** Dot plot summarizes cardiomyocyte CSA of healthy control (n=6), untreated BV173 (n=5) and DAS/VEN/DEX treated BV173 (n=5) hearts at 35 weeks FU after transplantation. ****p<0.0001 vs. Control, ^###’^p<0.0001 vs. leukemia. **(D-E)** Left ventricular cardiac output (LV CO) (D) and stroke volume (E) of age matched healthy controls (control) (n=5) or follow-up mice treated with DAS/VEN/DEX (n=4) 35 weeks after transplantation. Underlying data can be found in the file Supplement Figure 2-source data.

**Supplemental Figure 3:**
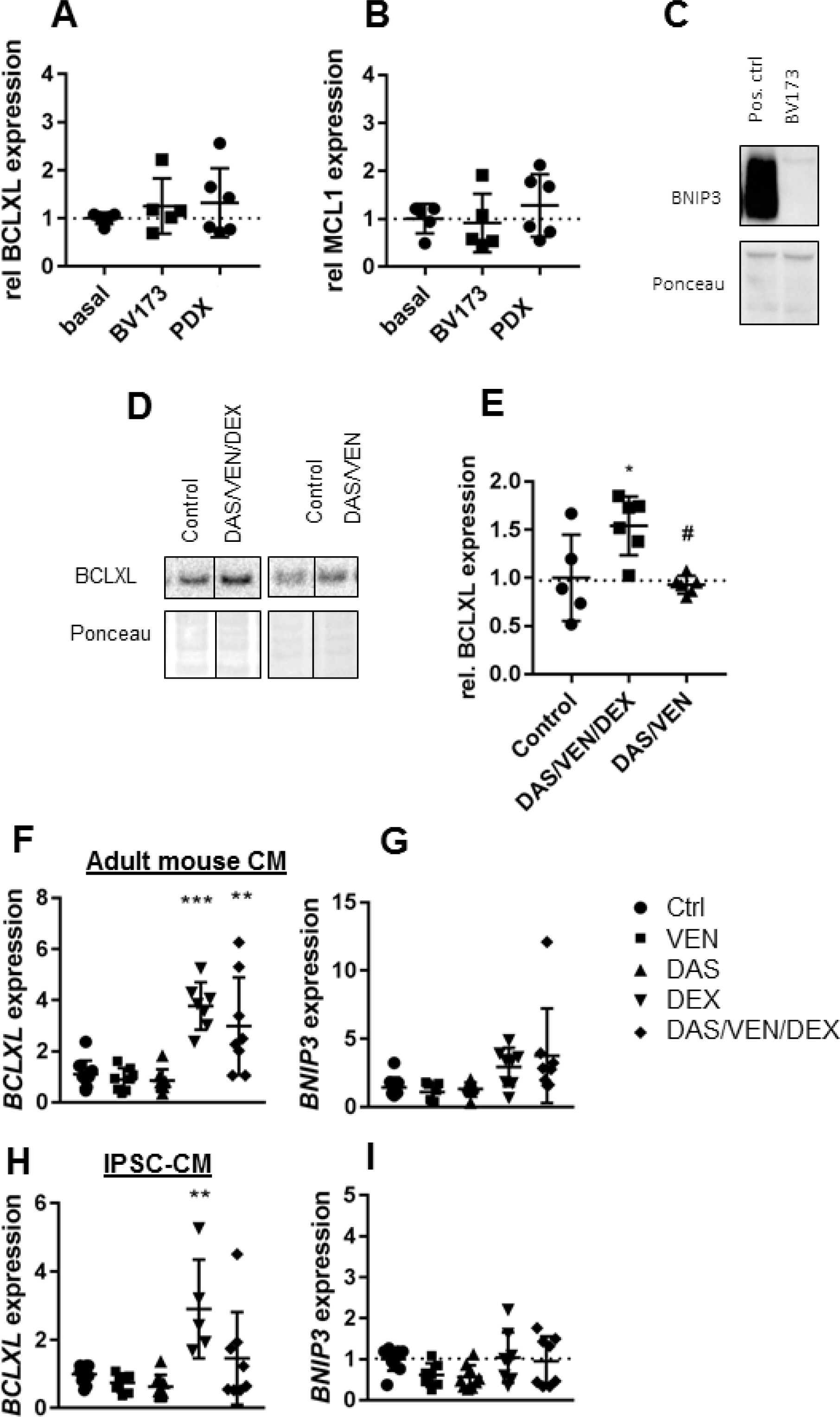
**(A-B)** Densitometric analysis of BCLXL (A) and MCL1 (B) expression in LV tissue of healthy NSG mice (n=5) or BV173-(n=5) and PDX-mice (n=6). Quantification was normalized to Ponceau loading control. **(C)** BNIP3 Immunoblot of BV173 cells. Ponceau served as loading control. **(D-E)** Immunoblot and densitometric quantification of LV tissue of NSG mice treated with vehicle (n=5), DAS/VEN/DEX (n=6) or DAS/VEN (n=6) for 4 weeks. *p<0.05 vs Control, #p<0.05 vs Leukemia. **(F-G)** BCLXL and BNIP3 expression of adult mouse CM treated with DAS, VEN, DEX or a combination. ***p<0.001, **p<0.01. **(H-I)** BCLXL and BNIP3 expression of iPS-CM treated with DAS, VEN, DEX or a combination. **p<0.01. Underlying data can be found in Supplement Figure 3-source data folder.

### Supplemental tables

Supplemental tables 1-4 and legends are provided in Supplementary file 1.

### Source data file legends

**Figure 1-source data.xlsx. Numerical raw data**. All numerical raw data of Figure 1 are combined in this single excel file.

**Figure 1H BNIP3-source data.scn**. Original file of the full raw unedited blot of Figure 1H BNIP3.

**Figure 1H BNIP3-source data.tif**. Uncropped blot image of Figure 1H BNIP3.

**Figure 1H Ponceau-source data.scn**. Original file of the full raw unedited blot of Figure 1H Ponceau.

**Figure 1H Ponceau-source data.tif**. Uncropped blot image of Figure 1H.

**Figure 1H-source data 2.pptx**. Uncropped blot image with the relevant bands labelled of Figure 1H.

**Figure 2-source data.xlsx. Numerical raw data**. All numerical raw data of Figure 2 are combined in this single excel file.

**Figure 2G LC3B-source data.scn**. Original file of the full raw unedited blot of Figure 2G LC3B.

**Figure 2G LC3B-source data.tif**. Uncropped blot image of Figure 2G LC3B.

**Figure 2G Ponceau-source data.tif**. Uncropped blot image of Figure 2G Ponceau.

**Figure 2G-source data 2.pptx**. Uncropped blot image with the relevant bands labelled of Figure 2G.

**Figure 2I BCLXL-source data.scn**. Original file of the full raw unedited blot of Figure 2I BCLXL.

**Figure 2I BCLXL-source data.tif**. Uncropped blot image of Figure 2I BCLXL.

**Figure 2I BNIP3-source data.scn**. Original file of the full raw unedited blot of Figure 2I BNIP3.

**Figure 2I BNIP3-source data.tif**. Uncropped blot image of Figure 2I BNIP3.

**Figure 2I GAPDH-source data.scn**. Original file of the full raw unedited blot of Figure 2I GAPDH.

**Figure 2I GAPDH-source data.tif**. Uncropped blot image of Figure 2I GAPDH.

**Figure 2I-source data 2.pptx**. Uncropped blot image with the relevant bands labelled of Figure 2G BCLXL, BNIP3 and GAPDH.

**Supplement Figure 1-source data.xlsx. Numerical raw data**. All numerical raw data of Supplemental Figure 1 are combined in this single excel file.

**Supplement Figure 1I BNIP3-source data.tif**. Uncropped blot image of Supplemental Figure 1I BNIP3.

**Supplement Figure 1I Ponceau-source data.tif**. Uncropped blot image of Supplemental Figure 1I Ponceau.

**Supplement Figure 1I-source data 2.pptx**. Uncropped blot image with the relevant bands labelled of Figure 1I BNIP3 and Ponceau.

**Supplement Figure 2-source data.xlsx. Numerical raw data**. All numerical raw data of Supplemental Figure 2 are combined in this single excel file.

**Supplement Figure 3-source data.xlsx. Numerical raw data**. All numerical raw data of Supplemental Figure 3 are combined in this single excel file.

**Supplement Figure 3C BNIP3-source data.scn**. Original file of the full raw unedited blot of Figure 3C BNIP3.

**Supplement Figure 3C BNIP3-source data.tif**. Uncropped blot image of Supplemental Figure 3C BNIP3.

**Supplement Figure 3C Ponceau-source data.tif**. Uncropped blot image of Supplemental Figure 3C Ponceau.

**Supplement Figure 3D BCLXL left-source data.scn**. Original file of the full raw unedited blot of Figure 3D BCLXL.

**Supplement Figure 3D BCLXL right-source data.scn**. Original file of the full raw unedited blot of Figure 3D BCLXL.

**Supplement Figure 3D BCLXL left-source data.tif**. Uncropped blot image of Supplemental Figure 3D BCLXL.

**Supplement Figure 3D BCLXL right-source data.tif**. Uncropped blot image of Supplemental Figure 3D BCLXL.

**Supplement Figure 3D Ponceau left-source data.scn**. Original file of the full raw unedited blot of Figure 3D Ponceau.

**Supplement Figure 3D Ponceau right-source data.scn**. Original file of the full raw unedited blot of Figure 3D Ponceau.

**Supplement Figure 3D Ponceau left-source data.tif**. Uncropped blot image of Supplemental Figure 3D Ponceau.

**Supplement Figure 3D Ponceau right-source data.tif**. Uncropped blot image of Supplemental Figure 3D Ponceau.

**Supplement Figure 3C-source data 2.pptx**. Uncropped blot image with the relevant bands labelled of Figure 3C BNIP3 and Ponceau.

**Supplement Figure 3D-source data 2.pptx**. Uncropped blot image with the relevant bands labelled of Figure 3D BCLXL, BNIP3 and Ponceau.

### Supplemental tables

**Suppl. Table 1:**
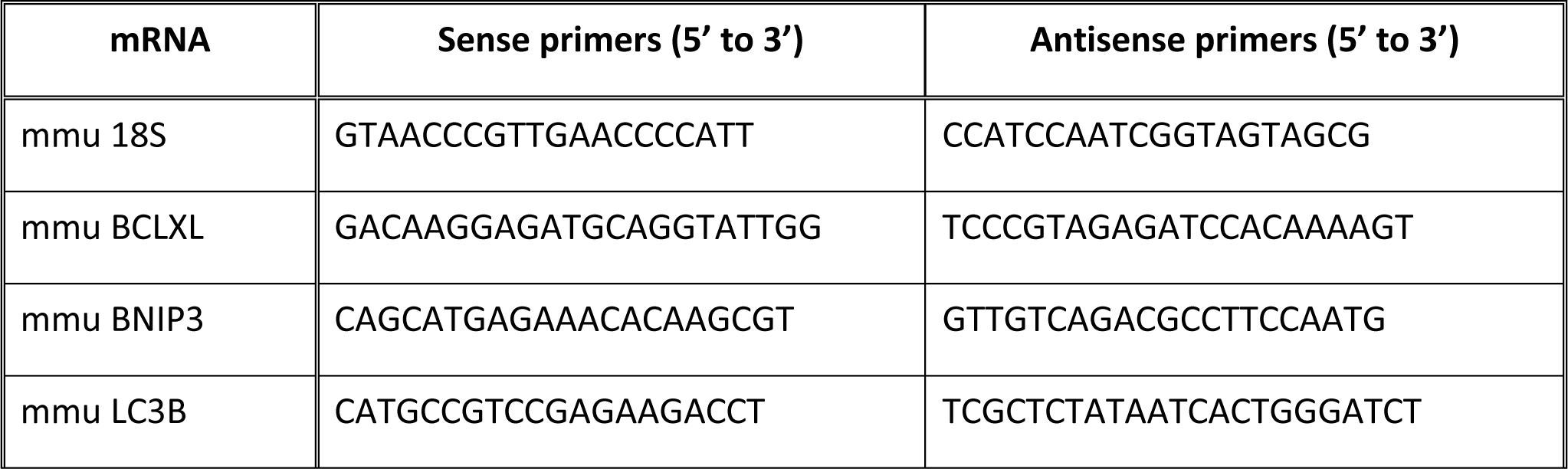
SYBR Green qRT-PCR Primer sequences.

**Supplement Table 1.**
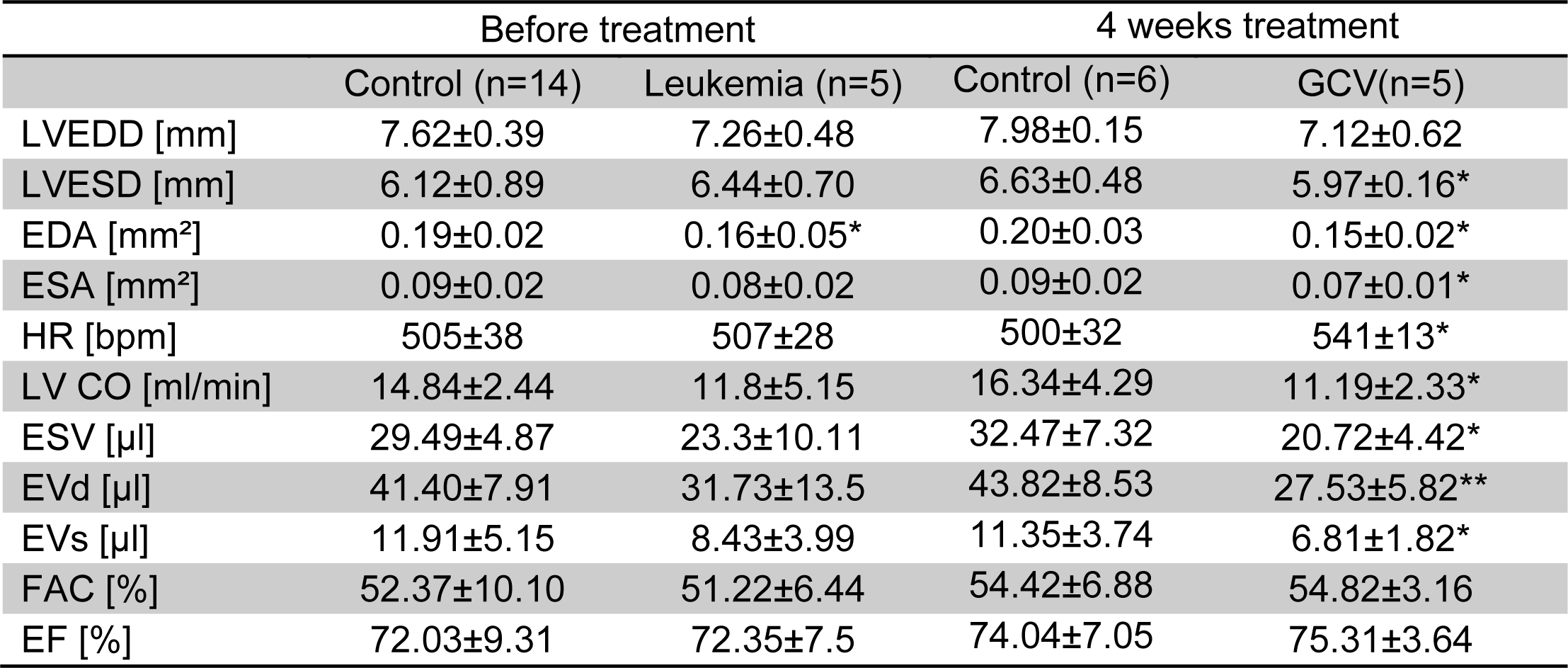
Cardiac function of NSG mice transplanted with BCR-ABL+ ALL. After confirmed state of advanced disease mice were treated with GCV for 4 weeks. Left ventricular end-diastolic diameter (LVEDD), left ventricular end-systolic diameter (LVESD), end-diastolic area (EDA), end-systolic area (ESA), heart rate (HR), left ventricular cardiac output (LV CO), endocardial stroke volume (EVS), endocardial diastolic volume (EVd), endocardial systolic volume (EVs), fractional area change (FAC), ejection fraction (EF); *p<0.05, **p<0.01 students t-test to corresponding aged matched healthy control group.

|  | Before treatment |  | 4 weeks treatment |  |
| --- | --- | --- | --- | --- |
|  | Control (n=14) | Leukemia (n=5) | Control (n=6) | GCV(n=5) |
| LVEDD [mm] | 7.62±0.39 | 7.26±0.48 | 7.98±0.15 | 7.12±0.62 |
| LVESD [mm] | 6.12±0.89 | 6.44±0.70 | 6.63±0.48 | 5.97±0.16* |
| EDA [mm<sup>2</sup>] | 0.19±0.02 | 0.16±0.05* | 0.20±0.03 | 0.15±0.02* |
| ESA [mm<sup>2</sup>] | 0.09±0.02 | 0.08±0.02 | 0.09±0.02 | 0.07±0.01* |
| HR [bpm] | 505±38 | 507±28 | 500±32 | 541±13* |
| LV CO [ml/min] | 14.84±2.44 | 11.8±5.15 | 16.34±4.29 | 11.19±2.33* |
| ESV [μl] | 29.49±4.87 | 23.3±10.11 | 32.47±7.32 | 20.72±4.42* |
| EVd [μl] | 41.40±7.91 | 31.73±13.5 | 43.82±8.53 | 27.53±5.82** |
| EVs [μl] | 11.91±5.15 | 8.43±3.99 | 11.35±3.74 | 6.81±1.82* |
| FAC [%] | 52.37±10.10 | 51.22±6.44 | 54.42±6.88 | 54.82±3.16 |
| EF [%] | 72.03±9.31 | 72.35±7.5 | 74.04±7.05 | 75.31±3.64 |

**Supplement Table 2.** Toxicity study 4 weeks of treatment in non-transplanted male NSG mice. Mice were treated with 75mg/kg/d GCV 5 days/week for 4 weeks. Left ventricular end-diastolic diameter (LVEDD), left ventricular end-systolic diameter (LVESD), end-diastolic area (EDA), end-systolic area (ESA), heart rate (HR), left ventricular cardiac output (LV CO), endocardial stroke volume (EVS), endocardial diastolic volume (EVd), endocardial systolic volume (EVs), fractional area change (FAC), ejection fraction (EF); no significance students t-test.

|  | Control (n=7) | GCV (n=6) |
| --- | --- | --- |
| LVEDD [mm] | 7.65±0.36 | 8.06±0.44 |
| LVESD [mm] | 6.70±0.26 | 6.92±0.34 |
| EDA [mm<sup>2</sup>] | 0.20±0.03 | 0.22±0.02 |
| ESA [mm<sup>2</sup>] | 0.10±0.01 | 0.12±0.01 |
| HR [bpm] | 504±35 | 528±34 |
| LV CO [ml/min] | 15.49±4.09 | 19.39±4.18 |
| ESV [μl] | 30.71±7.69 | 36.57±6.59 |
| EVd [μl] | 44.26±9.38 | 53.29±8.90 |
| EVs [μl] | 13.55±3.32 | 16.71±3.86 |
| FAC [%] | 48.18±6.27 | 48.17±5.16 |
| EF [%] | 69.10±6.78 | 68.66±4.93 |

**Supplement Table 3.** Morphometry and cardiac function of NSG mice transplanted with BCR-ABL+ ALL. After confirmed state of advanced disease mice were treated with DAS/VEN/DEX for 4 weeks. Body weight (BW), heart weight (HW), left ventricular end-diastolic diameter (LVEDD), left ventricular end-systolic diameter (LVESD), end-diastolic area (EDA), end-systolic area (ESA), heart rate (HR), left ventricular cardiac output (LV CO), endocardial stroke volume (EVS), endocardial diastolic volume (EVd), endocardial systolic volume (EVs), fractional area change (FAC), ejection fraction (EF); *p<0.05, **p<0.01, ***p<0.001 students t-test to corresponding aged matched healthy control group.

|  | Before treatment |  | 4 weeks treatment |  |
| --- | --- | --- | --- | --- |
|  | Control (n=14) | Leukemia (n=5) | Control (n=6) | DAS/VEN/DEX(n=5) |
| BW [g] | - | - | 24.62±1.34 | 19.20±3.15*** |
| HW [mg] | - | - | 102.50±7.94 | 98.86±7.23 |
| HW/BW [mg/g] | - | - | 4.17±0.35 | 5.14±0.32** |
| LVEDD [mm] | 7.62±0.39 | 7.63±0.15 | 7.98±0.15 | 7.82±0.31 |
| LVESD [mm] | 6.12±0.89 | 6.32±0.33 | 6.63±0.48 | 6.43±0.53 |
| EDA [mm<sup>2</sup>] | 0.19±0.02 | 0.18±0.02 | 0.20±0.03 | 0.18±0.02 |
| ESA [mm<sup>2</sup>] | 0.09±0.02 | 0.09±0.02 | 0.09±0.02 | 0.08±0.02 |
| HR [bpm] | 505±38 | 541±26 | 500±32 | 551±19* |
| LV CO [ml/min] | 14.84±2.44 | 11.87±2.52* | 16.34±4.29 | 13.69±2.28 |
| ESV [μl] | 29.49±4.87 | 22.07±5.23* | 32.47±7.32 | 24.91±4.38 |
| EVd [μl] | 41.40±7.91 | 35.46±9.1 | 43.82±8.53 | 34.25±6.69 |
| EVs [μl] | 11.91±5.15 | 13.39±5.07 | 11.35±3.74 | 9.34±2.95 |
| FAC [%] | 52.37±10.10 | 47.56±8.68 | 54.42±6.88 | 56.36±6.76 |
| EF [%] | 72.03±9.31 | 62.97±7.8 | 74.04±7.05 | 73.03±4.49 |

**Supplement Table 4.** Morphometry and cardiac function of long-term survivors. Follow-up (FU) of NSG mice transplanted with BV173 and treated six weeks with DAS/VEN/DEX 38 weeks after TX. Body weight (BW), heart weight (HW), left ventricular end-diastolic diameter (LVEDD), left ventricular end-systolic diameter (LVESD), end-diastolic area (EDA), end-systolic area (ESA), heart rate (HR), endocardial diastolic volume (EVd), endocardial systolic volume (EVs), fractional area change (FAC), ejection fraction (EF)no significance students t-test.

|  | Control (n=5) | FU DAS/VEN/DEX (n=4) |
| --- | --- | --- |
| BW [g] | 28.06±1.01 | 25.38±2.98 |
| HW [mg] | 102.46±6.83 | 90.18±3.56 |
| HW/BW [mg/g] | 3.65±0.15 | 3.56±0.08 |
| LVEDD [mm] | 7.34±0.21 | 7.53±0.75 |
| LVESD [mm] | 6.27±0.42 | 6.49±0.51 |
| EDA [mm<sup>2</sup>] | 0.19±0.01 | 0.19±0.03 |
| ESA [mm<sup>2</sup>] | 0.10±0.01 | 0.10±0.02 |
| HR [bpm] | 531±20 | 513.75±8.66 |
| EVd [μl] | 43.08±5.20 | 40.67±9.91 |
| EVs [μl] | 14.55±1.86 | 13.23±3.93 |
| FAC [%] | 46.88±6.89 | 47.12±4.70 |
| EF [%] | 65.63±7.19 | 67.51±4.69 |

